# Single-cell landscapes of long non-coding RNAs in early vascular endothelial development and hemogenic specification

**DOI:** 10.1101/2024.05.24.595647

**Authors:** Xupeng Chen, Xiaowei Ning, Chenguang Lu, Han He, Yingpeng Yao, Yanli Ni, Jie Zhou, Bing Liu, Siyuan Hou, Yu Lan, Zongcheng Li

## Abstract

Understanding the molecular regulation of arterial and hemogenic specification during early embryonic vascular development is crucial for guiding vascular and hematopoietic regeneration. Accumulating evidence emphasizes the role of long non-coding RNAs (lncRNAs) in cell fate decision. However, the dynamic expression and the potential roles of lncRNAs in early vascular development are still unknown. Here, we first constructed a single-cell landscape of lncRNA expression based on the deeply sequenced tag-based single-cell transcriptome data of early embryonic vascular endothelial cells (VECs). We revealed the contribution of lncRNAs to VEC heterogeneity and identified 295 lncRNAs with specific expression in eight VEC populations. Furthermore, we identified a series of lncRNAs potentially involved in regulating the two waves of arterial specification and hemogenic specification. We uncovered a transient downregulation of *H19* in the hemogenic endothelial population during endothelial-to-hematopoietic transition. Additionally, we constructed a transcription factor regulatory network composed of 287 regulons for early VEC development. We further revealed differential activation patterns of regulons and modules in the eight VEC populations, and predicted potential lncRNA-regulon regulatory network. Moreover, unsupervised analysis of the lncRNA expression profile revealed novel VEC subpopulations strongly associated with the maturation of VECs, suggesting the prominent roles of lncRNAs in endothelial maturation. In summary, our study fills the gap in understanding of lncRNA regulatory networks in early vascular development and provides insights into the fields of vascular and hematopoietic regeneration research.

## Introduction

Vascular endothelial cells (VECs) constitute the luminal layer of blood vessels to execute perfusion function, which is extremely pivotal throughout embryonic and adult life. During mammalian embryonic development, VECs are formed de novo from mesodermal precursors, assemble into a primordial vascular network via vasculogenesis. VECs then grow into new vessels from pre-existing ones via angiogenesis and specify into arterial and venous cell fates to form a hierarchically organized vascular network [1–5]. In addition, VECs play important perfusion independent roles. The hematopoietic stem cells (HSCs) are believed to be derived from hemogenic endothelial cells (HECs), a type of specialized embryonic VECs, via endothelial-to-hematopoietic transition [6–9]. In recent years, with the development of single-cell sequencing technology, a transcriptome landscape of early embryonic VEC development is constructed [10,11]. The dynamic spatiotemporal settling of arteriovenous and hemogenic fates of VECs in mid-gestational mouse embryos was unveiled [10–15]. Previously, we have uncovered two distinct arterial VEC types, the major artery VECs and arterial plexus VECs, and revealed two developmental paths of VECs from primordial VECs. One is towards mature major artery VECs, experiencing early arterial VECs and the subsequent maturing major artery VECs. The other one is towards early plexus VECs and further vein and venous plexus VECs, with the final fate conversion to become arterial plexus VECs [10]. In addition, we also witnessed the segregation of HSC-primed HECs from maturing major artery VECs along the major artery maturation path [11]. These findings provide a comprehensive understanding of the endothelial evolution and molecular program in arteriogenesis as well as in HSC-primed HEC specification, and provide a valuable database resource.

Long non-coding RNAs (lncRNAs) are a class of RNA molecules that are longer than 200 nucleotides and do not encode for proteins. LncRNAs have been found to be involved in a wide range of biological processes, including gene regulation, cell fate decision, chromatin modification, and epigenetic regulation [16–18]. Several lncRNAs have emerged as important regulators of vascular biology and HSC development. For example, lncRNAs may regulate endothelial dysfunction by modulating endothelial cell proliferation (e.g. *MALAT1*, *H19*), angiogenesis (e.g. *MEG3*, *MANTIS*) or vascular remodeling (e.g. *ANRIL*, *SMILR*, *SENCR*, *MYOSLID*) [19]. More than 150 unannotated lncRNA specifically enriched in adult HSCs have been identified and two lncRNAs have been proven to be functionally required in HSC self-renewal and differentiation [20]. The lncRNA *H19* has been reported to be pivotal for the endothelial-to-HSC transition that in trans regulates the promoter de-methylation of several critical hematopoietic transcription factors (TFs), including *Runx1* and *Spi1* [21]. However, the landscape of lncRNAs in early vascular developmental context, especially in the developmental path of two distinct arterial VEC types and during hemogenic fate selection, is still lacking.

Here, we integrated previously published single-cell RNA sequencing (scRNA-seq) data and constructed the lncRNA expression landscapes of VECs during early mammalian vascular development. We identified a series of important lncRNAs in both arteriovenous and hemogenic specification during early vascular development. Moreover, we revealed that lncRNAs could play prominent roles in endothelial cell maturation in both human and mouse. Our results fill an important gap in our understanding of the transcriptional regulatory networks that involve lncRNAs, and shed light on the studies in the vascular and HSC regeneration field. Furthermore, our study has revealed fresh avenues for capitalizing on the wealth of high-throughput tag-based scRNA-seq data currently accessible, including data generated by 10x Genomics, by examining lncRNAs from innovative perspectives.

## Results

### Abundant lncRNA expression in early VEC development uncovered by STRT-seq data

Our previous studies mainly focused on protein-coding genes, deciphering the heterogeneity in endothelial cells, widespread venous arterialization and hemogenic specification during early vascular development in mammals [10,11] (Figure S1). Meanwhile, a highly valuable scRNA-seq data has been generated for further mining. This scRNA-seq data had an extremely high sequencing depth, averaging three million reads per cell (about 30 times more than a regular transcriptome data by 10X Genomics), which was generated by using a modified STRT-seq method [22,23] with a 3’ tag-based library construction strategy similar to that of 10X Genomics. Here, we comprehensively quantified the expression profiles of lncRNAs in early mouse vascular development, based on the high-precision scRNA-seq data and a complete lncRNA gene annotation information from GENCODE [24] (Figure S1). High sequencing depth allows us to detect the expression of 6,722 lncRNA genes, accounting for approximately 30% of all expressed genes (Figure 1A). The expressed lncRNAs mainly consisted of antisense, lincRNA, and ‘to be experimentally confirmed’ (TEC), which together accounted for approximately 85% of all expressed lncRNAs (Figure 1A). At the single-cell level, on average, approximately 630 lncRNAs and 6,946 protein-coding genes (totally 7,576 genes) were expressed in VECs (Figure 1B). The proportion of lncRNAs in the expressed genes at the single-cell level (8.3%, 630/7,576) was much lower than their overall detection rate (29.5%) (Figure 1A), which may be due to lower expression levels and thus more dropout events of lncRNAs in single-cell sequencing (Figure 1C). Like protein-coding genes, the number of lncRNAs expressed varied in eight different embryo proper (EP) populations defined in our previous work [10] (Figure 1C). The number of expressed lncRNAs and protein-coding genes were relatively higher in primordial VEC population (corresponding to EP0) and early plexus VEC population (corresponding to EP1) and highest in HEC population (corresponding to EP4) (Figure 1B).

**Figure 1.**
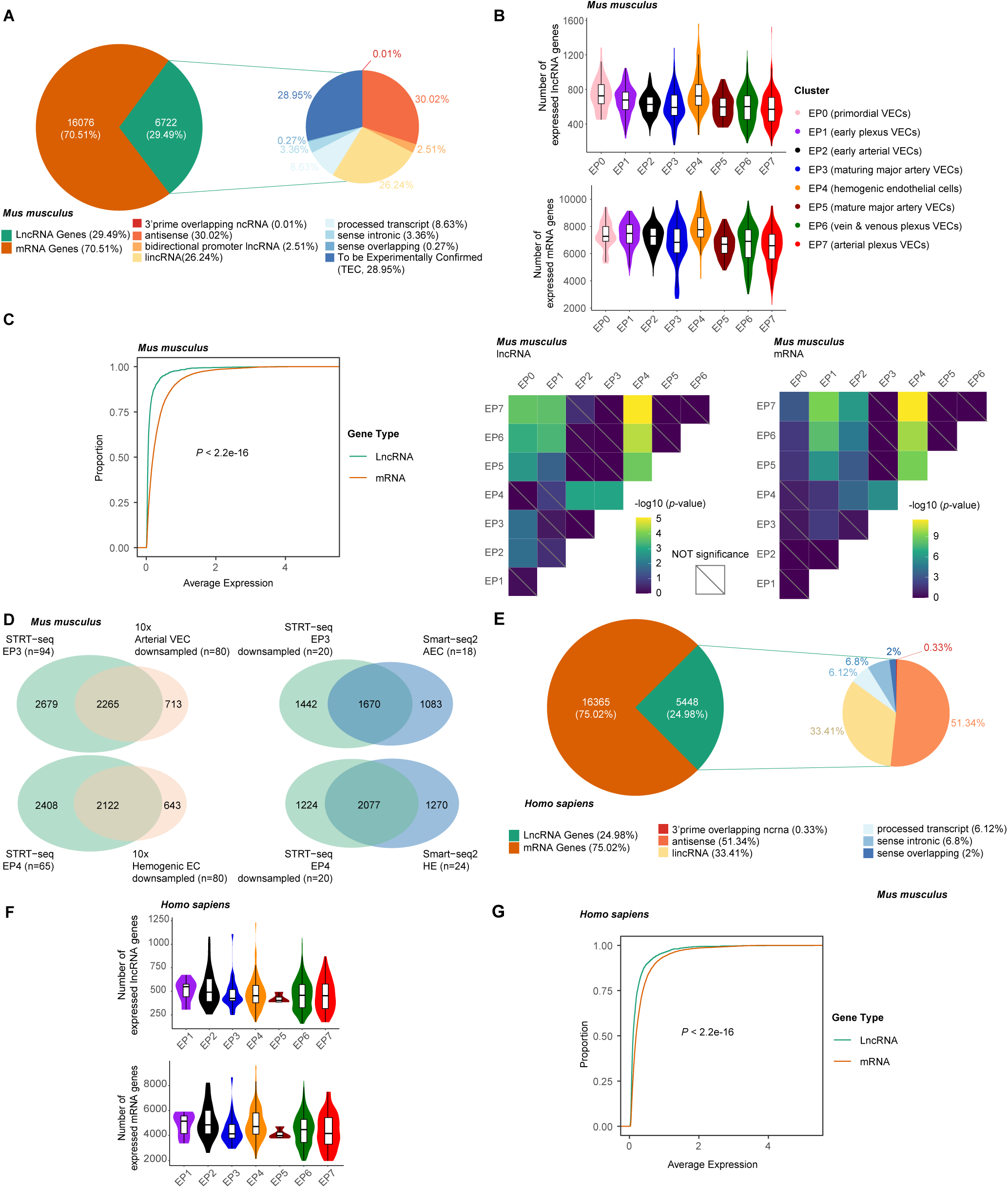
Abundant lncRNA expression in early VEC development uncovered by STRT-seq data. **A.** The number of protein-coding and lncRNA genes expressed during embryonic vascular development in *Mus musculus*. Only those genes expressed in more than three cells were classified as expressed genes for counting here. **B.** Number of expressed lncRNA and mRNA genes in individual cells for each cell type in *Mus musculus*. Adjusted p-values for differences between cell types are shown on the bottom. **C.** Cumulative distribution showing the expression level of lncRNA and mRNA genes in *Mus musculus*. The Kolmogorov-Smirnov test was used to determine the significance of differences. **D.** Venn diagram in left showing the overlap lncRNA genes detected by STRT-seq and 10x Genomics in AECs (top) and HECs (bottom), and in right showing the overlap lncRNA genes detected by STRT-seq and Smart-seq2 in AECs (top) and HECs (bottom). **E.** The number of protein-coding and lncRNA genes expressed during embryonic vascular development in *Homo sapiens*. Only those genes expressed in more than three cells were classified as expressed genes for counting here. **F.** Number of expressed lncRNA and mRNA genes in individual cells for each cell type in *Homo sapiens*. **G.** Cumulative distribution showing the expression level of lncRNA and mRNA genes in *Homo sapiens*. The Kolmogorov-Smirnov test was used to determine the significance of differences.

Additionally, we compared the number of lncRNAs detected in our STRT-seq to that in another tag-based sequencing data (e.g., 10x Genomics) and to that in the full-length sequencing data [25] (e.g., Smart-seq2) for the same VEC populations (i.e., EP3 vs. AEC vs. Arterial VEC and EP4 vs. HEC vs. Hemogenic EC) (Figure 1D). Our STRT-seq detected more lncRNAs than 10x Genomics and recovered most of the lncRNAs detected by 10x Genomics (approximately 76%). Our STRT-seq were comparable to the Smart-seq2 in the number of detected lncRNAs. Possibly due to differences in sequencing principles, each of the two strategies detects a biased variety of lncRNAs.

Similarly, we also quantified lncRNA expression in the modified STRT-seq data of human embryonic VEC development [10] and observed very similar results to those in mice. Briefly, 5,448 lncRNAs were detected, accounting for approximately 25% of all genes and mainly consisting of antisense and lincRNA (Figure 1E). On average, each cell expressed 471 lncRNAs and 4,544 protein-coding genes (Figure 1F). Compared to protein-coding genes, lncRNAs were also significantly enriched in lower expression level ranges (Figure 1G).

In summary, these results indicate that STRT-seq data with high sequencing depth can effectively and comprehensively capture lncRNA expression profiles in early vascular development.

### Differential expression of lncRNAs contributes to heterogeneity of VEC populations defined by protein-coding genes

We identified a total of 295 lncRNAs that were significantly differentially expressed between VEC populations (Figure 2A; Table S1). Using these differentially expressed lncRNAs as input, we performed dimensionality reduction analysis, and the UMAP plot clearly showed the relative concentration of cells in the same VEC population and the proximity relationships between VEC populations (Figure 2B). This result was also highly consistent with the dimensionality reduction results based on protein-coding genes [10,11], further demonstrating the significant contribution of these differentially expressed lncRNAs to the molecular heterogeneity among the eight VEC populations.

**Figure 2.**
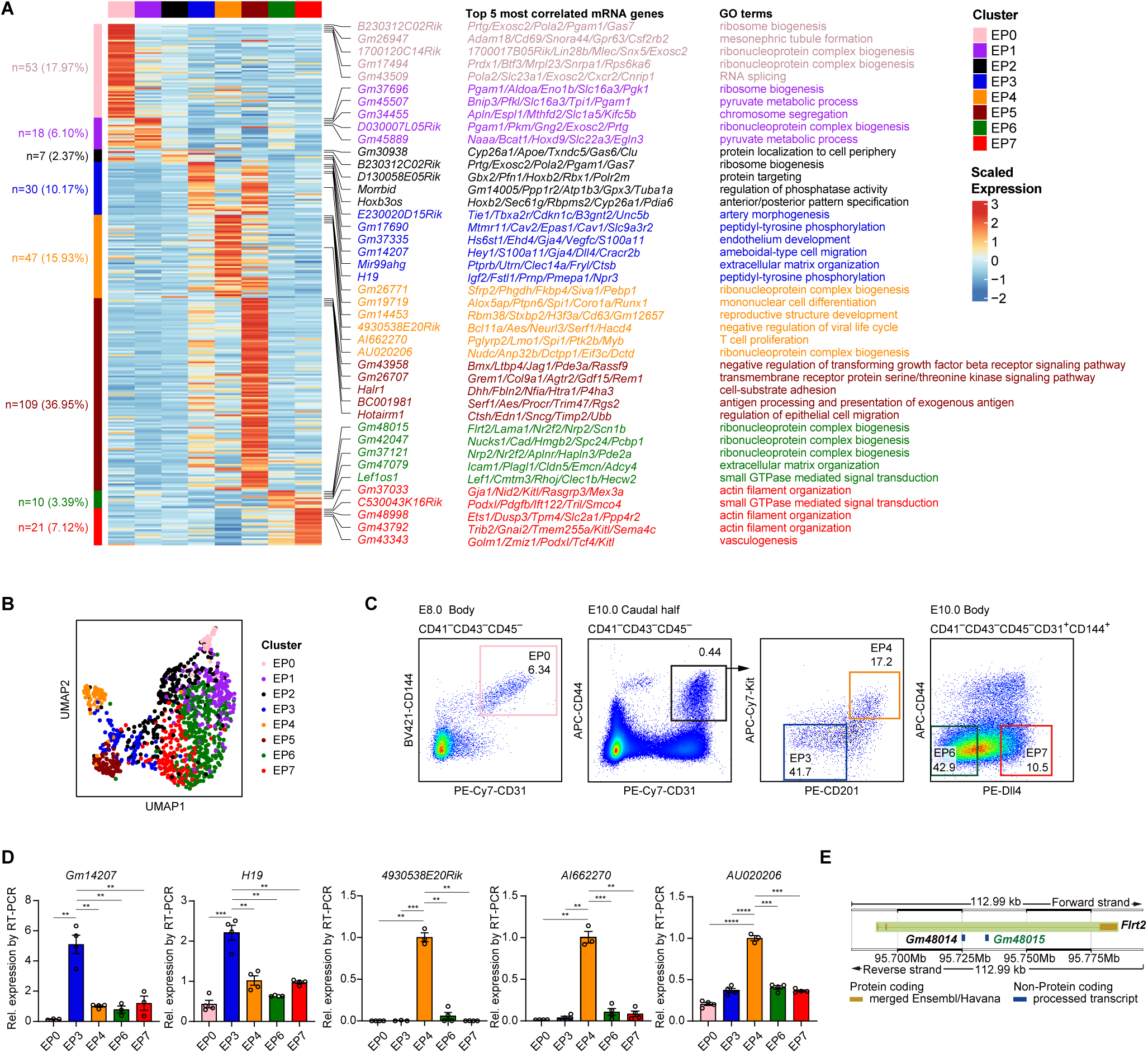
Differential expression of lncRNAs contributes to heterogeneity of VEC populations. **A.** Heatmap showing the average expression levels of DEGs in eight embryo proper VEC clusters. And the top 5 most correlated mRNA genes to the lncRNA genes and their enriched GO term, by using the top 100 most correlated protein-coding genes, were shown to the right. **B.** UMAP analysis of VECs during embryonic vascular development by using lncRNA DEGs as input. **C.** Representative FACS plots showing the sorting strategy of EP0, EP3, EP4, EP6, and EP7. **D.** Validation of two lncRNA DEGs for EP3 and three lncRNA DEGs for EP4 using the quantitative RT-PCR. Data are shown from at least three independent experiments. **E.** The genomic location of mRNA gene *Flrt2* and lncRNA genes *Gm48014* and *Gm48015*.

Given that the functions of the majority of lncRNA genes are unknown, we relied on the co-expressed protein-coding genes and their enriched biological processes to reflect the potential functional attributes of each lncRNA. For each population, we present the top five representative lncRNAs (i.e., top differentially expressed lncRNAs with enriched biological processes) (Figure 2A). Interestingly, these most correlated protein-coding genes and enriched biological processes varied among feature lncRNAs (Figure 2A), suggesting their potentially differentiated regulatory roles. Importantly, these predicted functions were consistent with the characteristics of the corresponding VEC populations. For example, *Gm14207* and *H19* were significantly higher in maturing arterial VECs (corresponding to EP3) (Figure 2A). *Gm14207* was significantly co-expressed with several important arterial feature genes, such as *Hey1*, *Gja4*, and *Dll4*, and enriched in the biological process of artery morphogenesis, suggesting the important regulatory role of *Gm14207* in arterial development. *H19* was enriched in the biological process of endothelial cell differentiation and was highly co-expressed with its imprinted gene partner *Igf2*, implying the important role of the *H19/Igf2* imprinting gene pair in early arterial endothelial development and differentiation. Among the representative lncRNAs of EP4, *Gm19719*, *4930538E20Rik*, *AI662270*, and *Gm9934* were highly co-expressed with hematopoiesis-related factors, such as *Spi1*, *Runx1*, *Neurl3* and *Adgrg1*. And *Gm14453* and *Gm9934* were significantly associated with definitive hemopoiesis and regulation of hemopoiesis (Figure 2A). It is worth noting that *AI662270* was also identified as a specific lncRNA of pre-HSC [21]. To further validate the accuracy of 3’ tag-based sequencing for lncRNA identification, we sorted the transcriptome-identified VEC populations using combinations of surface markers, as in our previous works [10,11], including primordial VECs (EP0) from E8.0 embryos, and maturing major artery VECs (EP3), HECs (EP4), vein & venous plexus VECs (EP6), and arterial plexus VECs (EP7) from E10.0 embryos (Figure 2C). Then we performed quantitative RT-PCR to validate the specific expression of two feature lncRNAs of EP3 (e.g., *Gm14207* and *H19*) and three feature lncRNAs of EP4 (e.g., *4930538E20Rik*, *AI662270* and *AU020206*) (Figure 2D; Tables S2 and 3). These results demonstrated the authenticity and expression specificity of these lncRNAs identified by 3’ tag-based sequencing. Additionally, these differentially expressed lncRNAs still contain a wealth of valuable information remaining to be explored and verified. For instance, we noticed that both *Gm48014* and *Gm48015* are two lncRNAs located in the intron of *Flrt2*, a mammalian conserved venous marker [10], but only the latter is significantly co-expressed with *Flrt2*, suggesting that *Gm48015* rather than *Gm48014* should be related to the transcriptional regulation of *Flrt2* in early VEC development (Figure 2A and E). In summary, our results reveal a series of lncRNAs with potentially important regulatory functions in early vascular development and hemogenic specification.

### lncRNAs regulate arteriovenous specification during early vascular development

Two waves of arterial specification, including the specification of the major artery (involving EP2, EP3, and EP5) and venous arterialization (involving EP1, EP6, and EP7), are the most prominent molecular events during early VEC development (Figure 3A). To reveal arteriovenous-associated lncRNAs, we identified the top 10 lncRNAs most strongly correlated with arterial and venous scores, respectively, such as *H19*, *Gm37335*, *Gm14207*, and *Malat1* for arterial features, and *D030007L05Rik*, *Gm42047*, *Snhg17*, and *Meg3* for venous features (Figure 3B). Previous studies have shown that lncRNAs can regulate vascular formation through mediating endothelial cell proliferation (e.g., *H19*, *Malat1*) or angiogenesis (e.g., *Meg3*) [19]. Our results showed that these lncRNAs could also help characterize the arterial and venous features of VEC and suggested their potential involvement in regulating arterial and venous specification.

**Figure 3.**
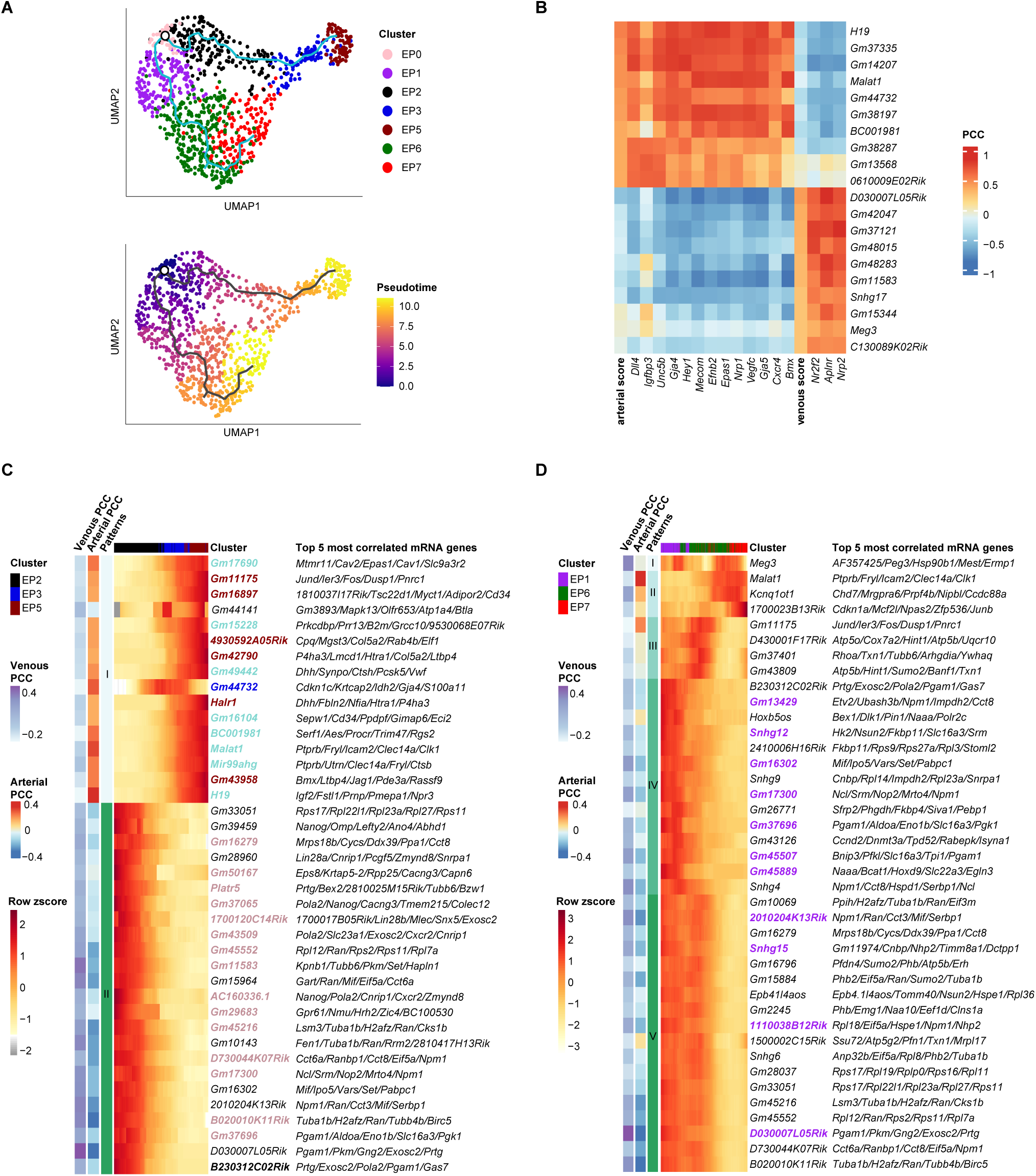
lncRNAs regulate arteriovenous specification during early vascular development. **A.** Developmental trajectory among embryo proper VEC clusters inferred by Monocle3 as previously published [10]. **B.** Heatmap showing the Pearson correlation coefficients of lncRNA genes with arterial score, venous score, and protein-coding genes used for calculating the arterial score or venous score. **C-D.** Heatmap showing the relative expression of the lncRNA genes along the development paths of two types of arterial VECs. Cells are ordered by pseudotime and genes are ordered by Patterns. And the correlation coefficient of genes and artery score or venous score were annotated to the left. The gene names with bold font were the DEGs of EC populations and the colors represented their belonging VEC clusters, except that the genes with fuchsia color are the DEGs of both EP3 and EP5.

Moreover, based on our previously established trajectories of the two-wave arterial development [10] (Figure 3A), we further analyzed the dynamic changes of lncRNAs along the arterial development pseudotime. In the path of major artery development, we identified 16 gradually up-regulated and 24 gradually down-regulated lncRNAs (Figure 3A and C). Almost all of the lncRNAs that were significantly positively correlated with development were also feature genes of EP3 and/or EP5 and were highly positively correlated with the arterial characteristic score (Figure 3C). Conversely, the lncRNAs that were significantly negatively correlated with development were mainly feature genes of early vascular endothelial EP0, such as *Platr5*, a pluripotency associated lncRNA [26]. These genes still have a high expression level in EP2 and gradually decrease as the artery matures (Figure 3C). In addition, some down-regulated lncRNAs were significantly positively correlated with the venous score, such as *D030007L05Rik* and *Gm11583*, which is consistent with the downregulation of venous features during artery development (Figure 3C). In the development path of venous arterialization, except for a few gradually up-regulated arterial feature lncRNAs (such as *Malat1*), most were gradually down-regulated venous feature lncRNA genes, including many differentially expressed lncRNAs of EP6 (Figure 3A and D). Additionally, multiple small nucleolar RNA (snoRNA) host genes (SNHGs), including *Snhg4/6/9/12/15*, were also involved in the down-regulated pattern with development (Figure 3D). The SNHG family has been found to play diverse roles in various cellular processes, including cell proliferation, differentiation, apoptosis, and migration [27]. And some of the most well-studied members of the SNHG family have been shown to be involved in various diseases, for example cancer progression and metastasis (e.g., SNHG1, SNHG5, and SNHG12), neurodegenerative diseases (e.g., SNHG6 and SNHG16), and cardiovascular disease (e.g., SNHG15) [27–29]. Our results suggest that the SNHG family lncRNAs may also play a potentially important role in early arterial specification of VECs.

### lncRNAs are involved in hemogenic endothelial specification

During early vascular development, in addition to the development and maturation of arterial VECs, there is a critical transient process named hemogenic endothelial specification. Gaining insight into the dynamic alterations of pivotal lncRNAs involved in the arterial VEC’s commitment to hematopoietic fate during endothelial to hematopoietic transition is a critical aspect in enhancing the regulatory network of hemogenic endothelial specification. To accomplish this, we utilized monocle 2 [30,31] to conduct pseudotemporal analysis and generated the developmental trajectories branching from EP3 to EP4 and to EP5 (Figure 4A). Then we identified lncRNAs that undergo dynamic changes along these two developmental paths (Figure 4B). Furthermore, these lncRNAs can be classified into four main patterns. Pattern 1 (P1) is only upregulated during HEC specification and represents hemogenic-specific lncRNAs including several EP4’s feature lncRNAs (Figure 4B). We constructed a co-expression network of these hemogenic-specific lncRNAs and their most correlated protein-coding genes, which contains a considerable number of EP4 high-expression protein-coding genes (Figure 4C). This suggests that these lncRNAs potentially play important roles in HEC specification. It is noteworthy that *AI662270* has been identified as a feature gene of pre-HSC [21]. In our study, we found that *AI662270*, together with *Gm19719*, was significantly linked with three hematopoietic-related TFs (including *Spi1*, *Runx1*, and *Myb*), indicating their possible cooperative regulation with key TFs (Figure 4C). P2 represents upregulated lncRNAs during arterial maturation, with multiple EP3/5 feature lncRNAs included (Figure 4B). P3 represents significantly downregulated lncRNAs in both HEC specification and arterial maturation, but more dramatic in the former (Figure 4B). The lncRNAs in P4 exhibit an entirely opposite pattern of changes between HEC specification and arterial maturation. For example, *H19* is downregulated during HEC specification but upregulated during arterial maturation (Figure 4B). We observed that previous research has reported upregulation of *H19* in pre-HSCs relative to VECs [21]. The present study suggests that HECs, as a preceding stage of pre-HSCs, undergo a transient downregulation of *H19* expression (Figure 4D). This result suggests a fine and complex regulatory mechanism of *H19* in the endothelial-to-hematopoietic transition process, and further exploration is required.

**Figure 4.**
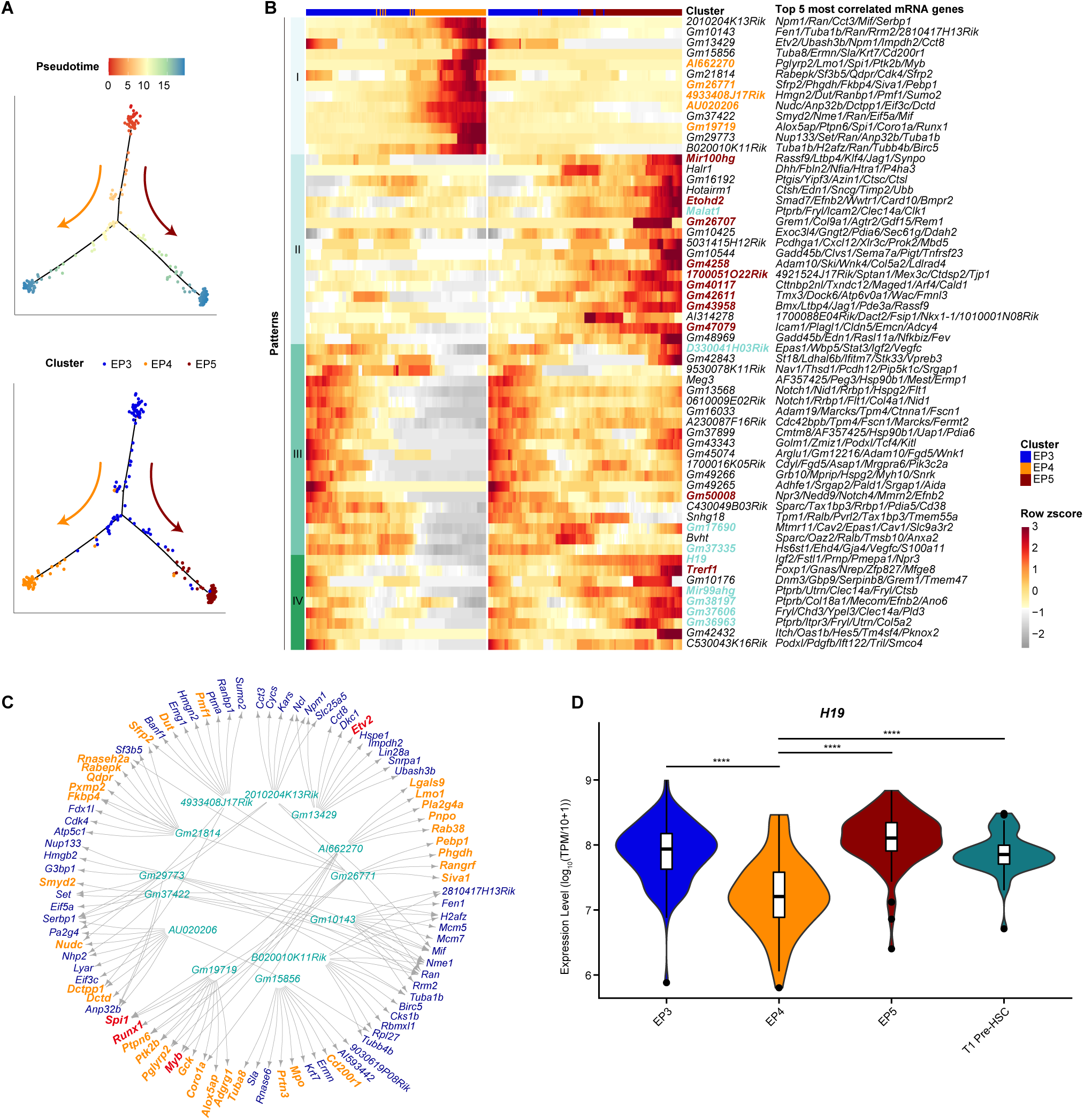
lncRNAs are involved in hemogenic endothelial specification. **A.** Pseudotemporal ordering of the cells in three major artery-related VEC clusters inferred by monocle 2, with clusters (top) and pseudotime (bottom) mapped to it. HEC specification and artery maturation directions are indicated as orange and deep red arrows, respectively. **B.** Heatmap showing the relative expression of the lncRNA genes with significantly dynamic changes along HECs specification and artery maturation paths. Cells are ordered by pseudotime and genes are ordered by patterns. The gene names with bold font were the DEGs of EC populations and their colors are consistent with their belonging VEC clusters, except that the genes with fuchsia color are the DEGs of both EP3 and EP5. **C.** Network diagram showing the relationship of lncRNA genes of patterns 1 and the top 10 most correlated protein-coding genes. The DEGs of EP4 were marked in yellow, and TFs were marked in red. **D.** Violin diagram shows the expression of H19 in the clusters as indicated.

### lncRNAs are associated with TF regulatory network in early vascular development

We employed SCENIC [32] to construct a regulatory network consisting of 287 regulons that instruct early vascular development. Through bi-directional unsupervised hierarchical clustering, we created a heatmap representation of the regulon activities, which effectively illustrated the comparable and distinctive regulon activation patterns among the eight VEC populations (Figure 5A). Furthermore, we calculated the connection specificity index (CSI) among regulons and categorized them into ten regulon modules based on the CSI matrix (Figure 5B). Interestingly, although there is no strong correlation among regulons within modules 1, 7, and 10, they are closely linked to other modules, such as module 1 to module 5, module 7 to modules 8 and 2, and module 10 to module 4 (Figure 5B and C). More importantly, the activity of these representative regulon modules shows differential activation among different populations and enriched in distinct biological processes (Figure 5C, Figure S2). Modules 1 and 5, relatively high activation in EP6 and EP7, were associated with small GTPase mediated signal transduction. Modules 2, 7 and 8, relatively high activation in EP3 and EP5, were associated with cell migration, morphogenesis and development. Similarly, Modules 4 and 10, high activation in EP4, were associated with ribonucleoprotein complex biogenesis, RNA processing, cell differentiation. Importantly, these results were also highly consistent with our previous observation of top enriched terms in DEGs (Figure 2A). Interestingly, module 10 also enriched in cell fate specification, which is in line with the endothelial-to-hematopoietic transition events experienced in EP4. In summary, these results reveal the TF regulatory network and its complex differential activation patterns, which may drive cluster-specific gene programs during early VEC development.

**Figure 5.**
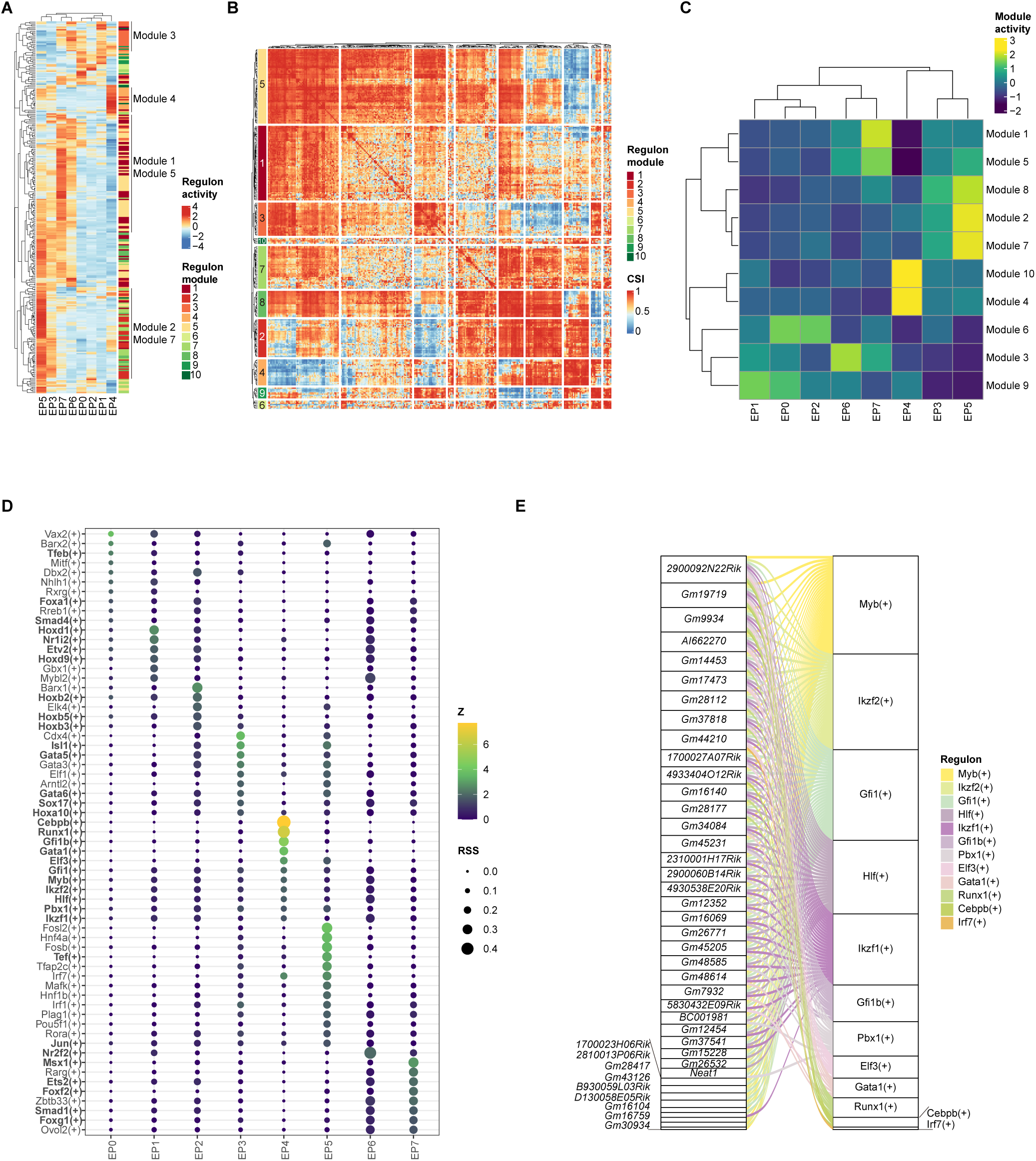
lncRNAs are associated with TF regulatory network in early vascular development. **A.** Heatmap showing the activity of all identified regulons in different VEC populations. The regulon modules (defined in panel B) are annotated to the right. **B.** Heatmap showing the CSI Scores between regulons. **C.** Heatmap showing the activity of all regulon modules in different VEC populations. **D.** Dot plot showing the regulons that specifically correlate with different VEC populations. The TFs which have been reported to be related with endothelial development or hematopoietic development were indicated in bold fonts. RSS, regulon specificity score. **E.** Alluvial diagram showing the potential regulatory network between lncRNAs and specific regulons in EP4 cluster.

We also utilized regulon specificity scoring to identify a series of VEC population-specific regulons (Figure 5D; Table S5). Of these 63 TFs, 35 have been reported in studies of endothelial development or hemogenic endothelial specialization. For instance, ETS factor *Etv2*, the master regulator of early vascular endothelial development[33,34], was significantly activated in early plexus VECs (i.e., EP1) (Figure 5D). Several members of the Hoxb family (e.g., *Hoxb2/3/5*), which were reported to be associated with endothelial cell activation and vessel formation [35], were specifically activated in EP2 (Figure 5D). *Sox17* and several Gata family factors (e.g., *Gata5/6*) were specifically activated during artery development and maturation (EP3 and EP5) (Figure 5D). *Sox17* is a key regulator of arterial identity [36], while *Gata5* and *Gata6* have been reported to play critical roles in the cardiovascular system [37,38]. The critical venous regulator *Nr2f2* was significantly activated in EP6, while *Msx1*, reported to be expressed in retina VECs at artery branching sites [39], was significantly activated in EP7 (Figure 5D). Nevertheless, the role of these reported endothelium-related TFs in arteriovenous specification of early vascular development remains to be determined. Meanwhile, those population-specific regulons with unknown functions in the endothelium were also warranted further investigation, especially those with stronger specificity (e.g., *Vax2* for EP0, *Barx1* for EP2, *Cdx4* for EP3, *Fosl2* and *Hnf4a* for EP5, and *Rarg* for EP5). (Figure 5D). Besides, several hemogenic-related TFs, including *Cebpb*, *Runx1*, *Gata1*, *Myb*, *Gfi1b* and *Gfi1*[21,40–42], were expectedly activated in HEC specification (EP4).

Next, we performed enrichment analysis to link lncRNAs with the specific regulons by calculating the significance of intersections between target genes in the regulon and top co-expressed protein-coding genes of lncRNAs (see Methods). We reasoned that those lncRNAs significantly associated with regulon may be potentially involved in the regulon regulatory network (e.g., transcriptional regulation of target genes). We further visualized the network of associations between lncRNAs and cluster-specific regulons (Figure 5E, Figure S3). Interestingly, some regulons showed strong correlation with multiple lncRNAs, including well-known early vascular development key regulator *Etv2*, venous specific TF *Nr2f2* and so on (Figure S3). Besides, tabking the EP4-specific lncRNA-regulon network for example, most of the regulons, including *Myb*, *Ikzf2*, *Gfi1*, *Hlf*, and *Ikzf1*, were associated with a greater number of lncRNAs, while *Runx1*, *Gata1* and *Cebpb* were associated with fewer lncRNAs (Figure 5E), indicating the differential involvement of lncRNAs in the regulatory network of the regulons of these TFs.

In summary, our findings elucidate the population-specific TF regulatory network and provide a list of candidate lncRNAs that potentially participate in the network.

### Expression of lncRNA distinguished novel VEC subpopulations related to endothelial maturation

We performed unsupervised dimensionality reduction and clustering analysis on the lncRNA expression profile and identified three major VEC populations (Figure 6A). EC3 included VECs belonging to EP3, EP5 and EP4, thus named as the major artery-related population. In line with the clear boundaries between distribution of EP3/4/5 in the UMAP plot, EC3 could be further subdivided as EP3/4/5 (Figure 6A, Figure S4A). However, the majorities of EP0, EP1 and EP2, about half of EP6 and a part of EP7 were mixed into EC1, while about half of EP6, most EP7 and a small part of EP1 and EP2 were together grouped into EC2 (Figure 6A and B). Obviously, newly identified EC1 and EC2 did not fit well with the previously defined EP clusters (Figure 6A). However, EC1 was enriched with VECs at earlier stages (e.g., mostly E8.0-E9.5), while EC2 was enriched with VECs at later stages (E9.5-E10.0) (Figure 6B). Moreover, we identified differentially expressed lncRNAs and protein-coding genes between EC1 and EC2 (Figure 6C, Figure S4C; Table S6). EC1 was mainly enriched in metabolism-related biological processes and highly expressed early endothelial-specific genes such as *Etv2* and *Lin28a*, while EC2 was enriched in endothelial cell development and differentiation (Figure 6C). Interestingly, those upregulated differentially expressed genes in EC2 exhibited relatively stronger expression in plexus VECs (EP6 and EP7) than early plexus or arterial VECs (EP1 or EP2, respectively) (Figure 6C, Figure S4C). Based on these observations, we speculate that EC1 and EC2 could distinguish the different differentiation state of VECs, and EC2 may represents more mature VECs. To validate this speculation, we employed CytoTRACE [43] to predict the differentiation/maturation state of VECs from all plexus VECs (EP1, EP6 and EP7). Strikingly, the change direction of CytoTRACE scores (i.e., maturity) almost perfectly matched the boundary between EC1 and EC2 but not the subtypes defined by protein-coding genes (Figure 6D). And EC1 and EC2 could distinguish VECs with different maturity levels independently of developmental stages and VEC subtypes (Figure 6E). These results suggest that the contribution of lncRNAs to VEC heterogeneity is predominantly in endothelial maturation during early embryonic developmental.

**Figure 6.**
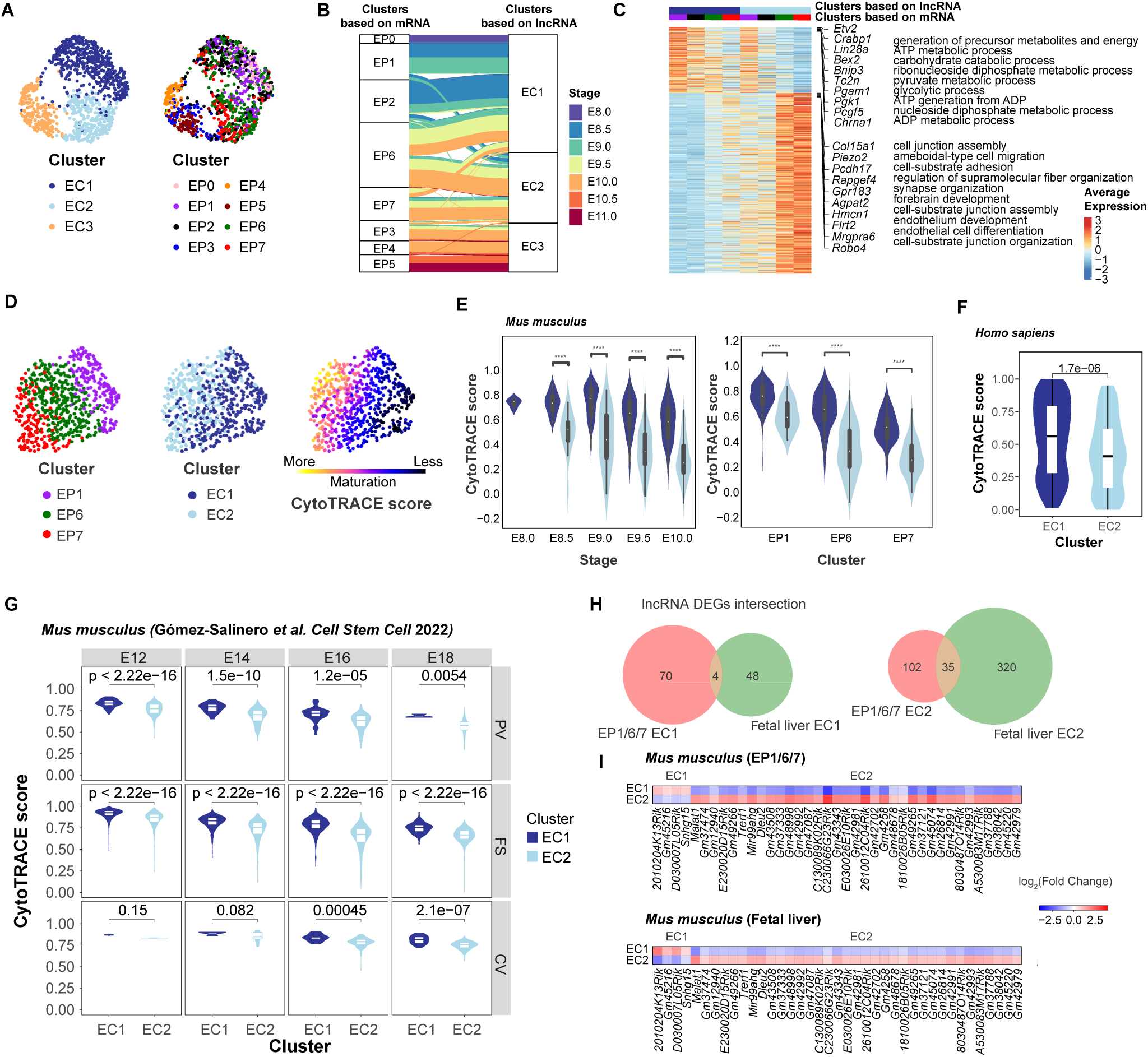
Expression of lncRNA distinguished novel VEC subpopulations. **A.** UMAP plots show the result of unsupervised clustering using lncRNA expression profiles(left), and embryo proper VEC clusters (right) of all VECs. **B.** The alluvial diagram shows the link between protein-coding gene-based clusters (EP0-7) lncRNA-based clusters (EC1-3) with link bands colored by stages. **C.** Heatmap showing the differentially expressed protein-coding genes between the EC1 and EC2 in the split VEC populations by cross-combination of EC1/2 and EP1/2/6/7. The top 10 DEGs and the most prominent GO enrichment terms of differential expression protein-coding genes are annotated to the right. D. UMAP plots show VEC clusters defined by our previous work (left), VEC clusters defined in this study based on lncRNA expression profiles (middle), and the CytoTRACE score (right) of EP1/6/7 populations. The UMAP dimension reduction was performed based on combined expression profiles of protein-coding genes and lncRNAs. E. Violin plots illustrate the differences in CytoTRACE scores of EC1/2 within the same developmental stage (left) and the same VEC clusters (right) in *Mus musculus*. F. Violin plots illustrate the differences in CytoTRACE scores of EC1/2 within the VECs of *Homo sapiens*. G. Violin plots illustrate the differences in CytoTRACE scores of EC1/2 within the same developmental stage and the same cell types in embryonic liver of *Mus musculus*. H. Venn diagrams illustrate the overlap (left) between DEGs of EC1 in EP1/6/7 and DEGs of EC1 in fetal liver, and the overlap (right) between DEGs of EC2 in EP1/6/7 and DEGs of EC2 in fetal liver. I. Heatmaps illustrate the differential expression fold changes of the overlapping DEGs between EC1 and EC2 in EP1/6/7 (top) and fetal liver (bottom).

Furthermore, we wondered whether this characteristic of lncRNAs is conserved in endothelial development of other scenarios (e.g., specific organs and different species). Here, we took mouse liver vascular development [44] and human early embryonic vascular development [10] for example. Surprisingly, unsupervised analyses on lncRNA expression profiles of these developing VECs uncovered highly consistent results with the observation above (Figure 6F and G, Figure S4D and E). Especially in liver vascular development, the identified clusters could distinguish VECs with different maturity levels independently of developmental stages and VEC subtypes defined based on protein-coding genes (Figure 6G; Table S7). Additionally, we identified 39 lncRNAs conservatively related to endothelial maturation in mouse, including *Snhg15*, *Malat1* and so on (Figure 6H and I), which warrant further research to confirm as endothelial maturation-related signature lncRNAs.

In conclusion, unlike protein-coding genes that primarily regulate arteriovenous specialization, lncRNAs may contribute to endothelial heterogeneity primarily by regulating endothelial cell maturation.

## Discussion

In this study, we found that rich lncRNA expression information can be mined from tag-based scRNA-seq data of early vascular endothelial development. In terms of detectable lncRNA quantity, tag-based data with high sequencing depth even exceeded full-length sequencing data. Further advanced analyses revealed that lncRNAs contributed to the heterogeneity of VECs and likely played an important regulatory role in key events, including arteriovenous specification, hemogenic specification and endothelial cell maturation, during early vascular development. It is worth noting that this study provides an example for the re-mining of high-throughput tag-based scRNA-seq data from the perspective of lncRNAs.

Based on supervised analyses, we revealed 295 differentially expressed lncRNAs, which contributed significantly to the heterogeneity of VEC populations, and a series of lncRNAs involved in arterial specification and hemogenic endothelial specification during early vascular development. We predicted the functions of lncRNAs by using co-expressed protein-coding genes. Moreover, we constructed the TF regulatory network instructing early VEC development, and identified a series of VEC population-specific regulons with a list of candidate lncRNAs that potentially participate in the regulon. Overall, this study provides valuable resources and insights into the regulatory roles of lncRNAs in early vascular development, especially regarding arterial, venous, and hemogenic specification. The identified feature lncRNAs and their predicted functions could serve as a foundation for further functional validation and exploration. Future studies could focus on investigating the molecular mechanisms underlying the regulatory roles of these lncRNAs and their potential applications in regenerative medicine and disease therapies.

Unexpectedly, unsupervised clustering based on lncRNA profiles revealed novel endothelial maturation associated VEC populations, which were independent of the arteriovenous associated EP clusters identified previously based on protein-coding genes. The findings highlight the importance of lncRNAs in regulation of endothelial cell development and maturation. Here, we only demonstrated that lncRNA expression profiles strongly associated with the maturation of VECs in early embryonic development of mouse and human, and mouse liver development. However, whether the dominant roles of lncRNA in other developmental scenarios of the endothelium are also conserved remains to be further verified.

The seemingly inconsistent results between supervised and unsupervised analyses, based on lncRNA expression profiles, actually reflect the heterogeneity of lncRNAs at different levels. Unsupervised analysis results reveal that the most highly variable lncRNAs are dominantly associated with endothelial maturation rather than endothelial arteriovenous specialization. Nonetheless, there is still a considerable amount of lncRNAs associated with endothelial arteriovenous specialization, which were uncovered by supervised analyses.

## Materials and Methods

### Mice

C57BL/6 mice were purchased from the Jackson Laboratory and housed in the Laboratory Animal Center of Academy of Military Medical Sciences in accordance with institutional guidelines. Mouse experiments were approved by the Animal Care and Use Committee of the Institute (IACUC-DWZX-2021-056). Embryos were staged by somite pair (sp) counting: E8.0, 1–7 sp and E10.0, 31–35 sp. Primary embryonic single-cell suspension was digested by 0.1% type I collagenase.

### Flow cytometry and antibodies

Cells were stained by the following antibodies: CD31 (102524, BioLegend, San Diego, CA), CD41 (553848, eBioscience, Waltham, MA), CD43 (553270, BD Biosciences, Bergen County, NJ), CD45 (11-0451-82, eBioscience, Waltham, MA), CD144 (138013, Biolegend, San Diego, CA), CD44 (17-0441-83, BioLegend, San Diego, CA), CD201 (12-2012-80, eBioscience, Waltham, MA), Kit (47-1171-82, eBioscience, Waltham, MA), and Dll4 (130807, BioLegend, San Diego, CA). 7-amino actinomycin D (7-AAD; 00-6993-50, eBioscience, Waltham, MA) was used to exclude dead cells. Cells were sorted by flow cytometers FACS Aria II (BD Biosciences, Bergen County, NJ). We sorted the transcriptome-identified VEC populations using combinations of surface markers, as in our previous works [10,11]: primordial VECs (CD41^-^CD43^-^CD45^-^ CD31^+^CD144^+^ for EP0) from E8.0 embryos, maturing major artery VECs (CD41^-^ CD43^-^CD45^-^CD31^+^CD44^+^Kit^-^CD201^-^ for EP3), HECs (CD41^-^CD43^-^CD45^-^ CD31^+^CD44^+^Kit^+^CD201^+^ for EP4), and plexus VECs (CD41^-^CD43^-^CD45^-^ CD31^+^CD144^+^CD44^-^Dll4^-^ and CD41^-^CD43^-^CD45^-^CD31^+^CD144^+^CD44^-^Dll4^+^ for EP6 and EP7, respectively) from E10.0 embryos.

### Quantitative RT-PCR

The cDNA preparation procedure was based on STRT with some modifications [22]. In brief, about 200 cells were sorted into 10 μL lysis buffer without ERCC. cDNAs were synthesized using sample-specific 25 nt oligo dT primer containing 8 nt barcode and TSO primer for template switching. After reverse transcription and second-strand cDNA synthesis, the cDNAs were amplified by 16 cycles of PCR using ISPCR primer and 3’P2 primer. Then each sample was purified with zymo kit and Agencourt AMPure XP beads (Beckman). Then Quantitative RT-PCR was carried out with SuperReal PreMix Plus SYBR Green (Tiangen) on a Roche Real-Time System (Roche 480). Results were processed by Microsoft Excel and then normalized to *Gapdh* transcripts’ expression. Fold differences in expression levels were calculated according to the 2^−ΔΔCT^ method. All primers used are listed in Table S2.

### Single-cell datasets selection and preprocessing

#### STRT-seq

In this study, single-cell RNA-seq data from *Mus musculus* and *Homo sapiens* embryos provided by S. Hou et al. [10] were utilized (Data was obtained from Gene Expression Omnibus with the accession number GSE94877). The data of *Mus musculus* embryos comprised 1,106 cells from 7 embryonic stages (E8.0 to E11.0). And the data of *Homo sapiens* embryos were generated from 4 Carnegie stages (CS10 to CS13). GENCODE project data (version M25 for *Mus musculus*, version 19 for *Homo sapiens*) [24] was employed to annotate 13,217 lncRNAs for *Mus musculus* and 13,853 lncRNAs for *Homo sapiens*. Quality control of the FASTQ reads was performed with Cutadapt (version 4.2) [45], aligned to the reference genome with Hisat2 (version 2.2.1) [46] and extracted unique reads with HTSeq (version 2.0.2) [47] based on cell barcodes and UMIs.

#### Smart-seq2

To evaluate the lncRNA genes detected distinguished between smart-seq2 sequencing and STRT-seq, the data published in F. Wang et al. [25] of AEC and HEC from *Mus musculus* were selected for analysis. Low-quality reads and adapters were filtered using Fastp (version 0.23.2) [48] with default parameters. The trimmed reads were aligned to the *Mus musculus* genome from GENCODE (version M25) using Hisat2 (version 2.2.1) [46] with default parameter. The aligned BAM files were sorted and indexed using SAMtools (version 1.6) [49]. Uniquely mapped reads were obtained using HTSeq (version 2.0.2) [47].

### Seurat analysis procedure for scRNA-seq datasets

Further analysis and exploration of single-cell RNA sequencing data were performed using the Seurat R package (version 4.1.1) [50]. This included identifying highly variable genes (HVGs) and differentially expressed genes (DEGs), as well as reducing dimensionality using PCA or UMAP with default parameters.

### Correlation analysis of EC populations

The correlation of different EC populations was calculate by using tl.dendrogram function in Scanpy python package [51] with default parameters.

### Differential expression analysis

Differential expression Genes (DEGs) were identified using FindMarkers or FindAllMarkers functions with default Wilcoxon rank sum test and only genes detected in a minimum fraction of 0.25 cells in either of the two populations were considered. Genes with fold-change ≥ 1.5 and adjusted *P* value ≤ 0.05 were selected as DEGs.

### Correlation analysis of lncRNA genes and mRNA genes

To address the issue of data sparsity affecting correlation analysis, we applied metacell construction instead of using single cells. Metacells are groups of similar cells that originate from the same biological sample and are aggregated using hdWGCNA (version 0.2.01) [52], a package that employs the k-Nearest Neighbors (KNN) algorithm to identify and combine cells with similar expression profiles. The resulting metacell gene expression matrix has a lower sparsity than the original expression matrix and is more suitable for correlation analysis. The MetacellsByGroups function was used with parameters “k=10, max_shared = 3, min_cells = 10, target_metacells = 18” to perform metacell construction. And then the metacell expression matrix was used to conduct a correlation analysis between lncRNA genes and mRNA genes. The Pearson correlation coefficient was calculated as an indicator of the linear relationship between lncRNA genes and mRNA genes. The lncRNA-mRNA pairs with a correlation coefficient of more than 0.3 and a *P* value of less than 0.05 were considered to be relevant. The network of correlation was drawn using igraph (version 1.3.4) R package [53].

### GO enrichment analysis

To predict the function of a lncRNA, the top 100 mRNA genes associated with the given lncRNA were selected for enrichment analysis. GO biological process enrichment analysis was performed using clusterProfiler [54].

### Constructing single-cell trajectories

The developmental trajectories of two types of arterial VEC populations were reconstructed using Monocle3 (version 1.0.0) [55] with the same process as the previous report [10]. The developmental trajectory of HEC was constructed using Monocle (version 2.22.0) [30,31]. For the Monocle analysis, UMI count data including mRNA and lncRNA features of given cell populations were used as input. The official vignette with recommended parameters was followed. HVGs were identified as genes with more than 1.5 times fitted dispersion evaluated using dispersionTable function. To reduce the influence of the cell cycle effect, only HVGs that were not intersected with genes included in the direct cell cycle GO term (GO:0007049) were retained as ordering genes for ordering cells.

### Arterial and venous feature score

A set of arteriovenous marker genes was selected based on previous knowledge or inference from the artery development pattern genes. These included 13 arterial genes (*Bmx*, *Cxcr4*, *Dll4*, *Efnb2*, *Epas1*, *Gja4*, *Gja5*, *Hey1*, *Igfbp3*, *Mecom*, *Nrp1*, *Unc5b*, and *Vegfc*) and 3 venous genes (*Aplnr*, *Nr2f2*, and *Nrp2*) [56–59] were used to perform the arteriovenous feature score. The specific calculation is consistent with the previous report [10].

### Dynamic changing genes along the development analysis

For the development path of two types of arterial VEC populations, pattern genes changing along the paths were identified by using graph_test function of Monocle3 with its Moran’s I test for each lncRNA gene. Genes with a *P* value ≤ 0.01 and the top 40 highest moran’s I score were selected. For the development paths of HEC and AEC, pattern genes changing along each development path were identified by using the BEAM function of Monocle. The top 60 significantly changed genes were retained for visualization in heatmaps for each development path. Pattern genes were clustered using the K-means method, and the number of clusters was determined by manually checking the heatmap results from a larger to a smaller number of clusters.

### Gene regulatory networks analysis

The standard pySCENIC pipeline [60] was followed to perform the analysis of regulon activity. The input to pySCENIC (version 0.12.1) [32] was the gene count matrix. The co-expressed gene network for each transcription factor was reconstructed by applying the GENIE3 algorithm analysis to the expression matrix. Then, the potential targets of transcription factors co-expression modules were further filtered with default parameters and used to build the regulons. The AUCell analyzed the regulon activity (area under the curve) and determined the active regulons with AUCell default threshold. To view how specific each predicted regulon is for each cell type, a regulon specificity score (RSS) was calculated based on the Jensen-Shannon divergence, a measure of the similarity between two probability distributions. For this calculation, each vector of binary regulon activity that overlaps with the assignment of cells to a specific cell type was used. The regulons that have an RSS > 0.05, meanwhile having a Z-score > 1.5, were filtered and considered as cluster specific regulons. The connection specificity index (CSI) is a measure of connectedness between different regulons. Regulons that share high CSI likely co-regulate downstream genes and are responsible for cell function together. Based on the AUCs matrix for all regulons, CSI scores for all regulon pairs were calculated. For identifying modules of regulons based on CSI, Hierarchical Clustering with default parameters was used on the regulon activity matrix of each cell.

### Identification of correlation between lncRNAs and regulons

Fisher’s exact test was used to investigate the relationship between lncRNAs and regulons. We conducted three parallel analyses by using top 100, 200 or 500 genes co-expressed with the lncRNA, respectively. The adjusted *P* value of less than 0.05 was considered to be significant. To get a more robust result, only lncRNAs with significance in all three results were retained as lncRNAs associated with the corresponding regulon.

### CytoTRACE score

To access the maturation of VECs, CytoTRACE scores were calculated by using CytoTRACE core in CellRank2 python package[61] with default parameters.

## Ethics statement

Mouse experiments were approved by the Animal Care and Use Committee of the Institute. (IACUC-DWZX-2021-056).

## Data availability

Processed data of scRNA-seq data are available download from Zenodo with the DOI: 10.5281/zenodo.11194329.

## Code availability

All data were analyzed with standard programs and packages, as detailed above. Code is available on request.

## CRediT author statement

Xupeng Chen: Methodology, Investigation, Software, Visualization. Xiaowei Ning: Methodology, Investigation, Validation. Chenguang Lu: Investigation, Validation. Han He: Software. Yingpeng Yao: Validation. Yanli Ni: Validation. Jie Zhou: Validation. Bing Liu: Conceptualization, Supervision. Siyuan Hou: Conceptualization, Writing - Original Draft. Yu Lan: Conceptualization, Writing - review & editing, Supervision. Zongcheng Li: Conceptualization, Writing - Original Draft, Supervision. All authors have read and approved the final Manuscript.

## Acknowledgments

This work was supported by the National Key R&D Program of China (2021YFA1100901, 2020YFA0112400, 2022YFA1103501, 2021YFA1100102, and 2022YFA1105700), the National Natural Science Foundation of China (82270118, 82000111, 82200121, 81890991, and 31930054), and the Program for Guangdong Introducing Innovative and Entrepreneurial Teams (2017ZT07S347).

## Supplementary materials

**Figure S1.**
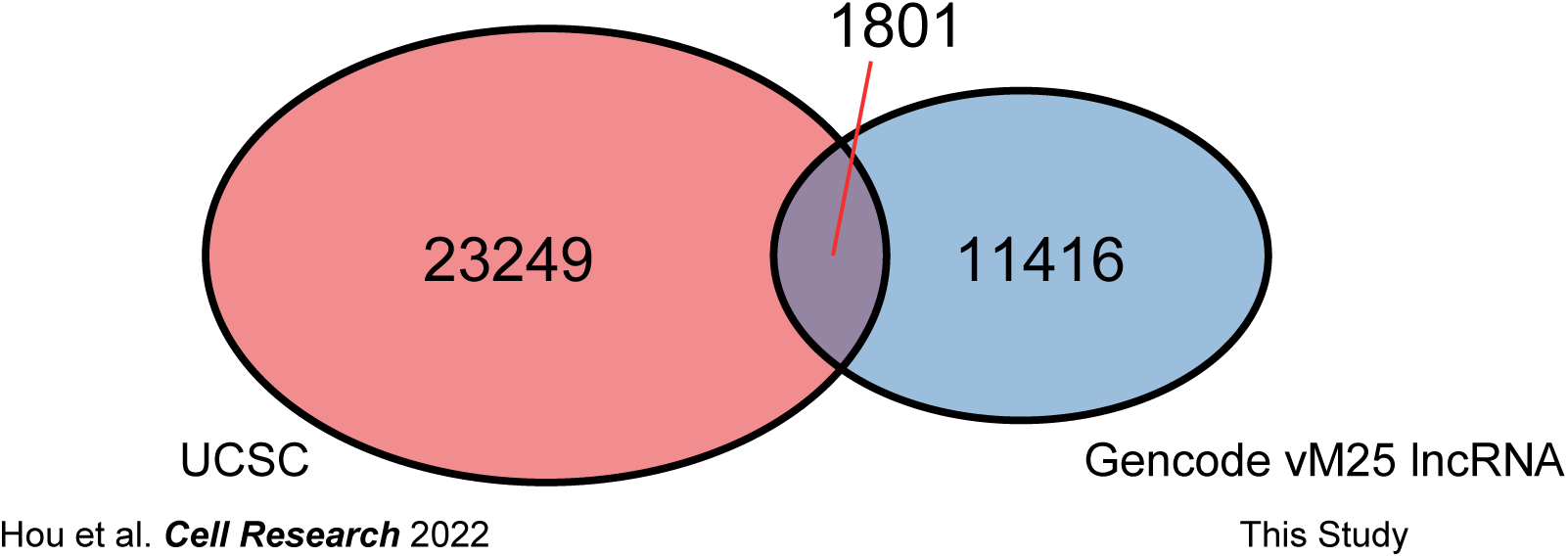
Comparison of gene annotations from previous work with the lncRNA gene annotations employed in this study.

**Figure S2.**
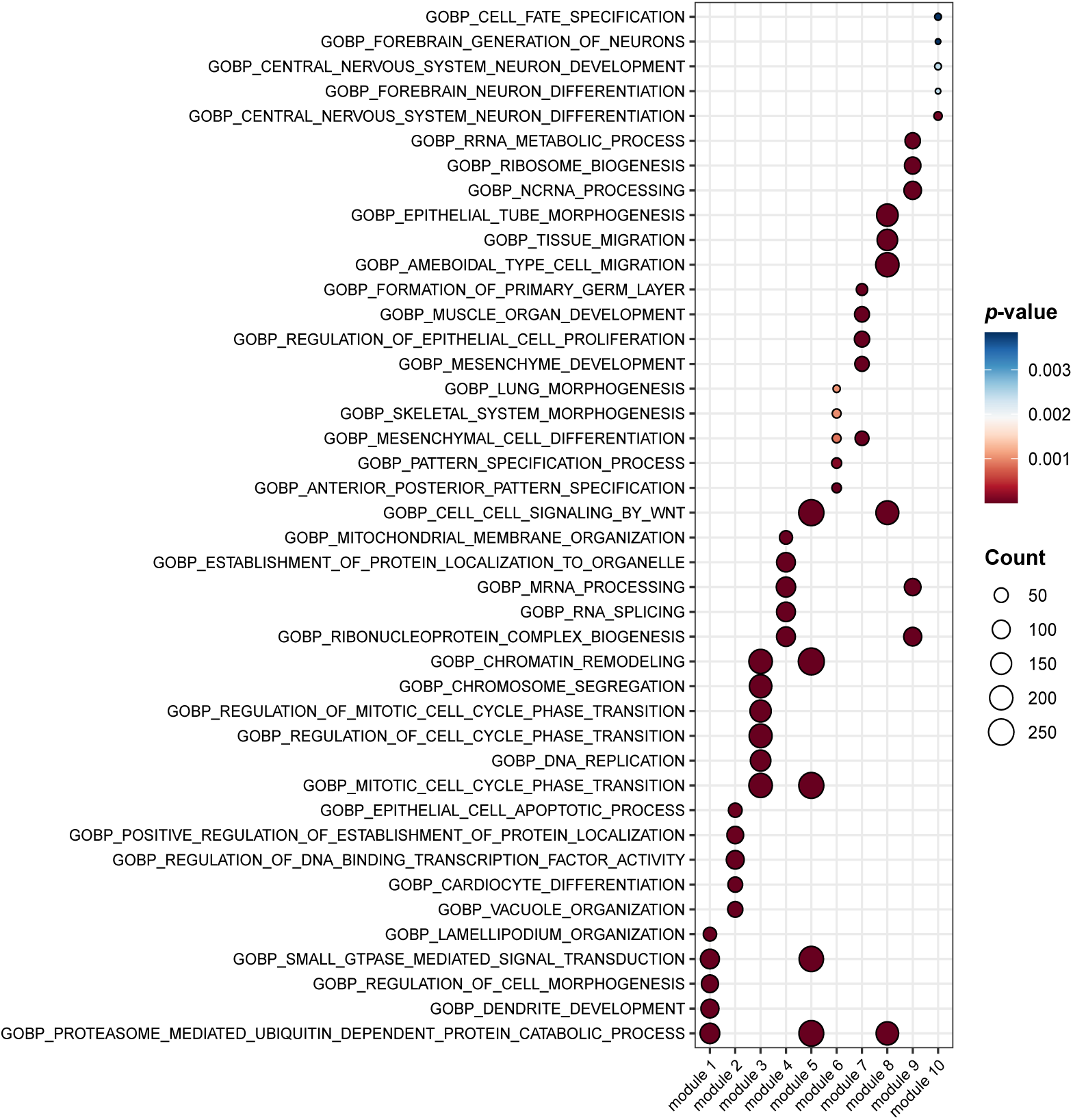
The GO enrichment terms of regulon modules.

**Figure S3.**
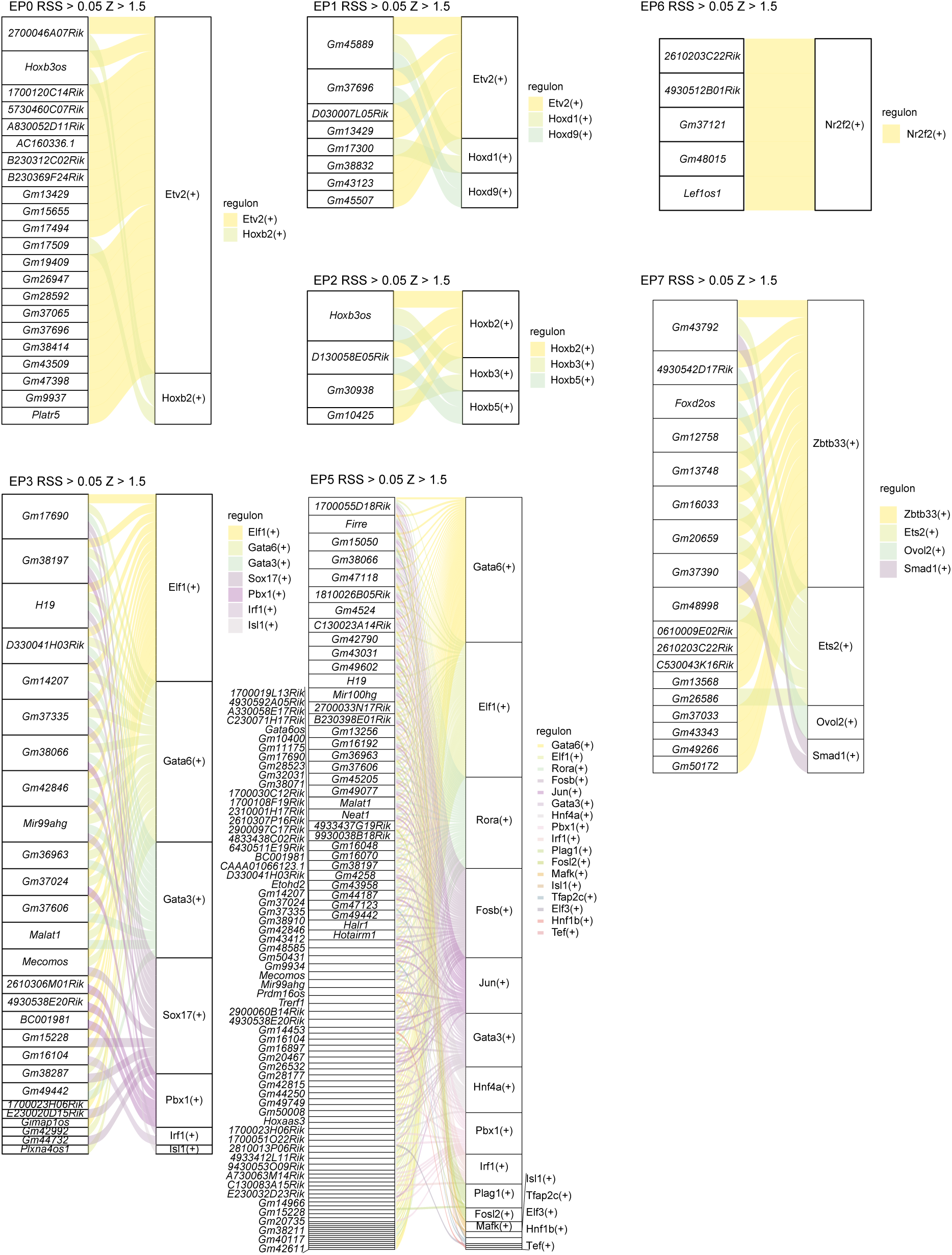
The potential regulatory network between lncRNAs and specific regulons in each cluster (except EP4 which has been shown in Fig. 5E).

**Figure S4.**
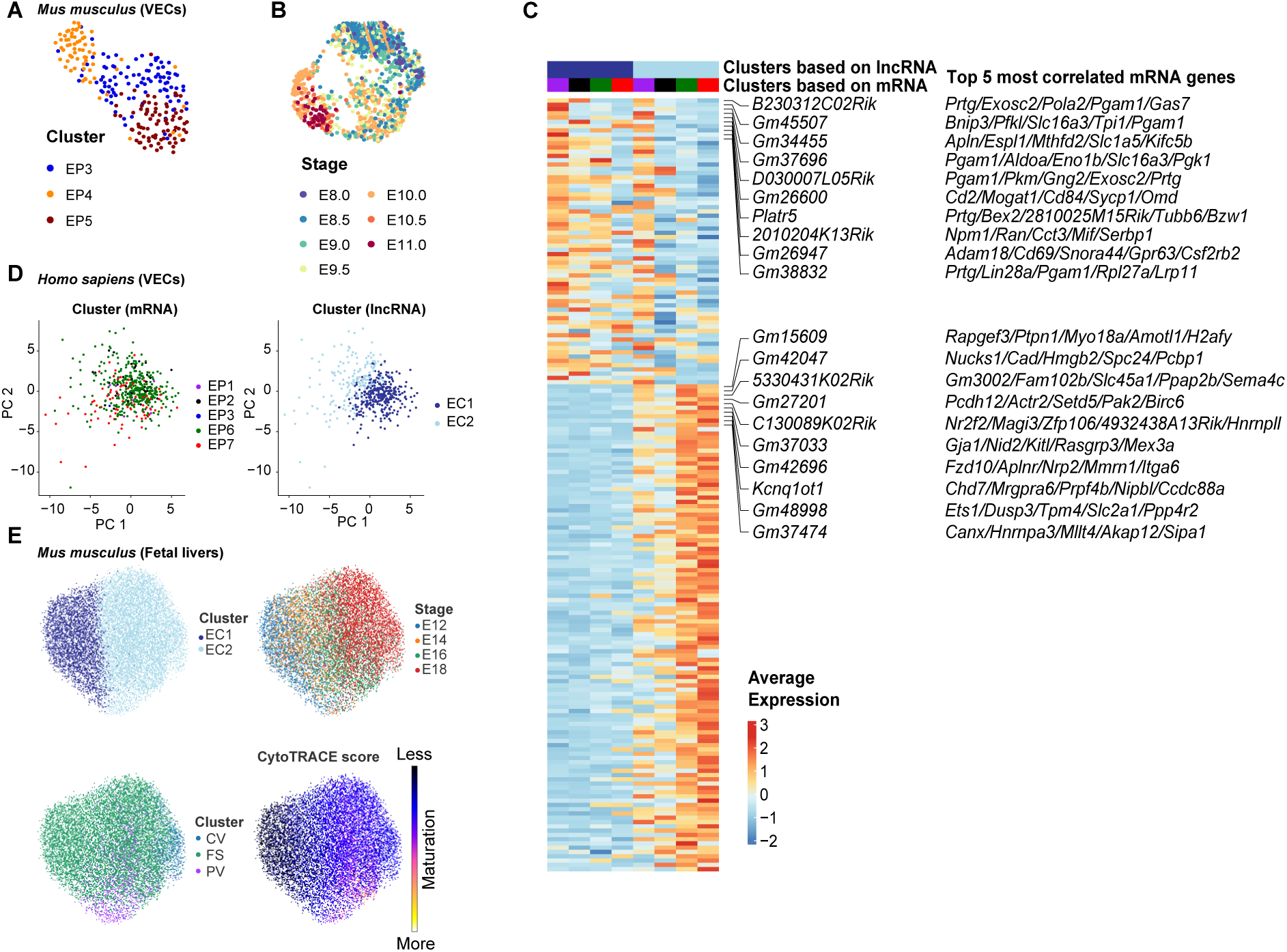
Expression of lncRNA distinguished novel VEC subpopulations. **A.** UMAP plots of VECs belonging to EC3 colored by EP3/4/5 populations. **B.** UMAP plots of all VECs colored by development stages. **C.** Heatmap showing the relative expression of differentially expressed lncRNAs between the EC1 and EC2 in the split VEC populations by cross-combination of EC1/2 and EP1/2/6/7. The mRNA genes significantly correlated to the lncRNAs are annotated to the right. **C.** PCA plots showing the result of unsupervised clustering using mRNA expression (left) and lncRNA expression profiles (right) of VECs in *Homo sapiens*. **D.** UMAP plots of VEC clusters identified based on lncRNA expression (top left), VEC clusters identified based on mRNA expression profiles (bottom left), developmental stage (top right), and CytoTRACE score (bottom right) in the fetal liver of *Mus musculus*. The UMAP dimension reduction was performed based on lncRNA expression profiles.

**Table S1. DEGs of VEC populations, related with Figure 2**.

**Table S2. q-PCR primers of candidate lncRNAs.**

**Table S3. Source data for q-PCR results of candidate lncRNAs, related with Figure 2**.

**Table S4. Regulon activity profiles in VEC clusters, related with Figure 5**.

**Table S5. Regulon module activities in VEC clusters, related with Figure 5**.

**Table S6. DEGs of novel VEC subpopulations defined by using lncRNAs, related with Figure 6**.

**Table S7 DEGs of novel EC subpopulations of fetal liver defined by using lncRNAs, related with Figure 6**.

## References

[1] Adams RH, Alitalo K. Molecular regulation of angiogenesis and lymphangiogenesis. Nat Rev Mol Cell Biol 2007;8:464–78.

[2] Herbert SP, Stainier DYR. Molecular control of endothelial cell behaviour during blood vessel morphogenesis. Nat Rev Mol Cell Biol 2011;12:551–64.

[3] Potente M, Mäkinen T. Vascular heterogeneity and specialization in development and disease. Nat Rev Mol Cell Biol 2017;18:477–94.

[4] Red-Horse K, Siekmann AF. Veins and Arteries Build Hierarchical Branching Patterns Differently: Bottom-Up versus Top-Down. BioEssays 2019;41:1800198.

[5] Trimm E, Red-Horse K. Vascular endothelial cell development and diversity. Nat Rev Cardiol 2023;20:197–210.

[6] Dzierzak E, Bigas A. Blood Development: Hematopoietic Stem Cell Dependence and Independence. Cell Stem Cell 2018;22:639–51.

[7] Kissa K, Herbomel P. Blood stem cells emerge from aortic endothelium by a novel type of cell transition. Nature 2010;464:112–5.

[8] Boisset JC, van Cappellen W, Andrieu-Soler C, Galjart N, Dzierzak E, Robin C. In vivo imaging of haematopoietic cells emerging from the mouse aortic endothelium. Nature 2010;464:116–20.

[9] Yokomizo T, Dzierzak E. Three-dimensional cartography of hematopoietic clusters in the vasculature of whole mouse embryos. Development 2010;137:3651–61.

[10] Hou S, Li Z, Dong J, Gao Y, Chang Z, Ding X, et al. Heterogeneity in endothelial cells and widespread venous arterialization during early vascular development in mammals. Cell Res 2022;32:333–48.

[11] Hou S, Li Z, Zheng X, Gao Y, Dong J, Ni Y, et al. Embryonic endothelial evolution towards first hematopoietic stem cells revealed by single-cell transcriptomic and functional analyses. Cell Res 2020;30:376–92.

[12] Zhu Q, Gao P, Tober J, Bennett L, Chen C, Uzun Y, et al. Developmental trajectory of prehematopoietic stem cell formation from endothelium. Blood 2020;136:845–56.

[13] Zhou F, Li X, Wang W, Zhu P, Zhou J, He W, et al. Tracing haematopoietic stem cell formation at single-cell resolution. Nature 2016;533:487–92.

[14] Baron CS, Kester L, Klaus A, Boisset JC, Thambyrajah R, Yvernogeau L, et al. Single-cell transcriptomics reveal the dynamic of haematopoietic stem cell production in the aorta. Nat Commun 2018;9:2517.

[15] Fadlullah MZH, Neo WH, Lie-a-ling M, Thambyrajah R, Patel R, Mevel R, et al. Murine AGM single-cell profiling identifies a continuum of hemogenic endothelium differentiation marked by ACE. Blood 2022;139:343–56.

[16] Flynn RA, Chang HY. Long noncoding RNAs in cell-fate programming and reprogramming. Cell Stem Cell 2014;14:752–61.

[17] Kallen AN, Zhou XB, Xu J, Qiao C, Ma J, Yan L, et al. The imprinted H19 lncRNA antagonizes let-7 microRNAs. Mol Cell 2013;52:101–12.

[18] Wang P, Xue Y, Han Y, Lin L, Wu C, Xu S, et al. The STAT3-binding long noncoding RNA lnc-DC controls human dendritic cell differentiation. Science 2014;344:310–3.

[19] Simion V, Haemmig S, Feinberg MW. LncRNAs in vascular biology and disease. Vascul Pharmacol 2019;114:145–56.

[20] Luo M, Jeong M, Sun D, Park HJ, Rodriguez BAT, Xia Z, et al. Long Non-Coding RNAs Control Hematopoietic Stem Cell Function. Cell Stem Cell 2015;16:426–38.

[21] Zhou J, Xu J, Zhang L, Liu S, Ma Y, Wen X, et al. Combined Single-Cell Profiling of lncRNAs and Functional Screening Reveals that H19 Is Pivotal for Embryonic Hematopoietic Stem Cell Development. Cell Stem Cell 2019;24:285–298.e5.

[22] Picelli S, Björklund ÅK, Faridani OR, Sagasser S, Winberg G, Sandberg R. Smart-seq2 for sensitive full-length transcriptome profiling in single cells. Nat Methods 2013;10:1096–8.

[23] Picelli S, Faridani OR, Björklund ÅK, Winberg G, Sagasser S, Sandberg R. Full-length RNA-seq from single cells using Smart-seq2. Nat Protoc 2014;9:171–81.

[24] A F, M D, Am F, R J, I J, J L, et al. GENCODE reference annotation for the human and mouse genomes. Nucleic Acids Research 2019;47.

[25] Wang F, Tan P, Zhang P, Ren Y, Zhou J, Li Y, et al. Single-cell architecture and functional requirement of alternative splicing during hematopoietic stem cell formation. Sci Adv 2022;8:eabg5369.

[26] Bergmann JH, Li J, Eckersley-Maslin MA, Rigo F, Freier SM, Spector DL. Regulation of the ESC transcriptome by nuclear long noncoding RNAs. Genome Res 2015;25:1336–46.

[27] Shi J, Ding W, Lu H. Identification of Long Non-Coding RNA SNHG Family as Promising Prognostic Biomarkers in Acute Myeloid Leukemia. Onco Targets Ther 2020;13:8441–50.

[28] Shuai Y, Ma Z, Lu J, Feng J. LncRNA SNHG15: A new budding star in human cancers. Cell Proliferation 2020;53:e12716.

[29] Zimta A-A, Tigu AB, Braicu C, Stefan C, Ionescu C, Berindan-Neagoe I. An Emerging Class of Long Non-coding RNA With Oncogenic Role Arises From the snoRNA Host Genes. Frontiers in Oncology 2020;10.

[30] Qiu X, Mao Q, Tang Y, Wang L, Chawla R, Pliner HA, et al. Reversed graph embedding resolves complex single-cell trajectories. Nat Methods 2017;14:979–82.

[31] Qiu X, Hill A, Packer J, Lin D, Ma Y-A, Trapnell C. Single-cell mRNA quantification and differential analysis with Census. Nat Methods 2017;14:309–15.

[32] Van de Sande B, Flerin C, Davie K, De Waegeneer M, Hulselmans G, Aibar S, et al. A scalable SCENIC workflow for single-cell gene regulatory network analysis. Nat Protoc 2020;15:2247–76.

[33] Palikuqi B, Nguyen D-HT, Li G, Schreiner R, Pellegata AF, Liu Y, et al. Adaptable haemodynamic endothelial cells for organogenesis and tumorigenesis. Nature 2020;585:426–32.

[34] Kim TM, Lee RH, Kim MS, Lewis CA, Park C. ETV2/ER71, the key factor leading the paths to vascular regeneration and angiogenic reprogramming. Stem Cell Research & Therapy 2023;14:41.

[35] Nova-Lampeti E, Aguilera V, Oporto K, Guzmán P, Ormazábal V, Zúñiga F, et al. Hox Genes in Adult Tissues and Their Role in Endothelial Cell Differentiation and Angiogenesis. Endothelial Dysfunction - Old Concepts and New Challenges, IntechOpen; 2018.

[36] Corada M, Orsenigo F, Morini MF, Pitulescu ME, Bhat G, Nyqvist D, et al. Sox17 is indispensable for acquisition and maintenance of arterial identity. Nat Commun 2013;4:2609.

[37] Tremblay M, Sanchez-Ferras O, Bouchard M. GATA transcription factors in development and disease. Development 2018;145:dev164384.

[38] Reiter JF, Alexander J, Rodaway A, Yelon D, Patient R, Holder N, et al. Gata5 is required for the development of the heart and endoderm in zebrafish. Genes Dev 1999;13:2983–95.

[39] Lopes M, Goupille O, Saint Cloment C, Robert B. Msx1 is expressed in retina endothelial cells at artery branching sites. Biology Open 2012;1:376–84.

[40] Fadlullah MZH, Neo WH, Lie-a-ling M, Thambyrajah R, Patel R, Mevel R, et al. Murine AGM single-cell profiling identifies a continuum of hemogenic endothelium differentiation marked by ACE. Blood 2022;139:343–56.

[41] Zeng Y, He J, Bai Z, Li Z, Gong Y, Liu C, et al. Tracing the first hematopoietic stem cell generation in human embryo by single-cell RNA sequencing. Cell Res 2019;29:881–94.

[42] Sato A, Kamio N, Yokota A, Hayashi Y, Tamura A, Miura Y, et al. C/EBPβ isoforms sequentially regulate regenerating mouse hematopoietic stem/progenitor cells. Blood Advances 2020;4:3343–56.

[43] Gulati GS, Sikandar SS, Wesche DJ, Manjunath A, Bharadwaj A, Berger MJ, et al. Single-cell transcriptional diversity is a hallmark of developmental potential. Science 2020;367:405–11.

[44] Gómez-Salinero JM, Izzo F, Lin Y, Houghton S, Itkin T, Geng F, et al. Specification of fetal liver endothelial progenitors to functional zonated adult sinusoids requires c-Maf induction. Cell Stem Cell 2022;29:593–609.e7.

[45] Martin M. Cutadapt removes adapter sequences from high-throughput sequencing reads. EMBnetJournal 2011;17:10–2.

[46] Kim D, Paggi JM, Park C, Bennett C, Salzberg SL. Graph-based genome alignment and genotyping with HISAT2 and HISAT-genotype. Nat Biotechnol 2019;37:907–15.

[47] Putri GH, Anders S, Pyl PT, Pimanda JE, Zanini F. Analysing high-throughput sequencing data in Python with HTSeq 2.0. Bioinformatics 2022;38:2943–5.

[48] Chen S, Zhou Y, Chen Y, Gu J. fastp: an ultra-fast all-in-one FASTQ preprocessor. Bioinformatics 2018;34:i884–90.

[49] Danecek P, Bonfield JK, Liddle J, Marshall J, Ohan V, Pollard MO, et al. Twelve years of SAMtools and BCFtools. GigaScience 2021;10:giab008.

[50] Hao Y, Hao S, Andersen-Nissen E, Mauck WM, Zheng S, Butler A, et al. Integrated analysis of multimodal single-cell data. Cell 2021;184:3573–3587.e29.

[51] Wolf FA, Angerer P, Theis FJ. SCANPY: large-scale single-cell gene expression data analysis. Genome Biology 2018;19:15.

[52] Morabito S, Reese F, Rahimzadeh N, Miyoshi E, Swarup V. High dimensional co-expression networks enable discovery of transcriptomic drivers in complex biological systems 2022:2022.09.22.509094.

[53] Csardi G, Nepusz T. The Igraph Software Package for Complex Network Research. InterJournal 2005;Complex Systems:1695.

[54] Wu T, Hu E, Xu S, Chen M, Guo P, Dai Z, et al. clusterProfiler 4.0: A universal enrichment tool for interpreting omics data. The Innovation 2021;2:100141.

[55] Cao J, Spielmann M, Qiu X, Huang X, Ibrahim DM, Hill AJ, et al. The single-cell transcriptional landscape of mammalian organogenesis. Nature 2019;566:496–502.

[56] Su T, Stanley G, Sinha R, D’Amato G, Das S, Rhee S, et al. Single-cell analysis of early progenitor cells that build coronary arteries. Nature 2018;559:356–62.

[57] Simons M, Eichmann A. Molecular Controls of Arterial Morphogenesis. Circulation Research 2015;116:1712–24.

[58] Chong DC, Koo Y, Xu K, Fu S, Cleaver O. Stepwise arteriovenous fate acquisition during mammalian vasculogenesis. Developmental Dynamics 2011;240:2153–65.

[59] Corada M, Morini MF, Dejana E. Signaling Pathways in the Specification of Arteries and Veins. Arteriosclerosis, Thrombosis, and Vascular Biology 2014;34:2372–7.

[60] Aerts’ Lab. Tutorial — pySCENIC latest documentation. Available from: https://pyscenic.readthedocs.io/en/latest/tutorial.html (accessed March 1, 2023).

[61] Lange M, Bergen V, Klein M, Setty M, Reuter B, Bakhti M, et al. CellRank for directed single-cell fate mapping. Nat Methods 2022;19:159–70.

