## Supplementary material for "Single-cell landscapes of long non-coding RNAs in early vascular endothelial development and hemogenic specification": Table S2

| Table S2 q-PCR primers of candiated lncRNAs | |
| --- | --- |
| LncRNA Gene | Sequence |
| *Gapdh-F* | *TCAACGACCACTTTGTCAAGCTCA* |
| *Gapdh-R* | *GCTGGTGGTCCAGGGGTCTTACT* |
| *H19-F* | *TCACTGAAGGCGAGGATGAC* |
| *H19-R* | *CCAGAGAGCAGCAGAGAAGT* |
| *GM14207-F* | *TTCCTTGGGCCACTTGAGAG* |
| *GM14207-R* | *TTCACGCTCTGCTCTTTCCC* |
| *AI662270-F* | *CCTCGGAGATGAAAGATGGACC* |
| *AI662270-R* | *CAAACTAACAGGCAGCCACGA* |
| *AU020206-F* | *AGCAGATGGAGGTGTTGTGT* |
| *AU020206-R* | *TCAAAGACCCTGAGATGGCC* |
| *4930538E20Rik-F* | *CACACACAGCCTTCGCTAT* |
| *4930538E20Rik-R* | *CGTGATTACAGTAGGTGCCC* |
