## Supplementary material for "Single-cell landscapes of long non-coding RNAs in early vascular endothelial development and hemogenic specification": Table S3

| Table S3 Source data for q-PCR results of candidate lncRNAs, related with Figure 2 | | | | | |
| --- | --- | --- | --- | --- | --- |
| Gene | Cluster | Experiment 1 | Experiment 2 | Experiment 3 | Experiment 4 |
| *AI662270* | EP0 | 0.013570961 | 0.014850617 | 0.014646164 | 0.012188519 |
|  | EP3 | 0.037725051 | 0.034955582 | 0.005160286 | 0.081427881 |
|  | EP4 | 1.142082339 | 0.91806402 | 0.953739165 |  |
|  | EP6 | 0.211197793 | 0.07113319 | 0.127331976 | 0.022046091 |
|  | EP7 | 0.09451366 | 0.041569427 | 0.027425594 | 0.174544484 |
| *AU020206* | EP0 | 0.243444813 | 0.199805287 | 0.224792634 | 0.155681203 |
|  | EP3 | 0.431271016 | 0.313528102 | 0.387338461 | 0.34547822 |
|  | EP4 | 1.008119503 | 0.943874313 | 1.050930065 |  |
|  | EP6 | 0.431271016 | 0.340721919 | 0.435778422 | 0.413702811 |
|  | EP7 | 0.346677633 | 0.360149215 | 0.395477265 | 0.357661483 |
| *4930538E20Rik* | EP0 | 0.000414885 | 0.000218512 | 0.000599065 | 0.000189568 |
|  | EP3 | 0.000438541 | 0.000419 | 0.007894152 |  |
|  | EP4 | 0.979420298 | 0.923382311 | 1.105730653 |  |
|  | EP6 | 0.174948233 | 0.001386 | 0.065607293 | 0.002093308 |
|  | EP7 | 0.000312252 | 0.001381068 | 0.000139253 | 0.000425074 |
| *H19* | EP0 | 0.709561678 | 0.335643126 | 0.392292049 | 0.292194312 |
|  | EP3 | 2.505328877 | 2.234574276 | 2.419988178 | 1.675974269 |
|  | EP4 | 1.257013375 | 0.784584098 | 0.864537231 | 1.172834949 |
|  | EP6 | 0.630688704 | 0.65747138 | 0.624165274 | 0.626332219 |
|  | EP7 | 1.031683179 | 0.989656656 | 0.989656656 | 0.882702996 |
| *Gm14207* | EP0 | 0.178469568 | 0.099355938 | 0.159182098 |  |
|  | EP3 | 3.467149391 | 6.314856718 | 4.937394768 | 5.671577248 |
|  | EP4 | 1.085793686 | 1.070845244 | 1.056102601 | 0.814366432 |
|  | EP6 | 1.192302244 | 0.773112356 | 0.385218795 |  |
|  | EP7 | 1.984464838 | 1.277877905 | 0.353247193 |  |
