## Supplementary material for "Single-cell landscapes of long non-coding RNAs in early vascular endothelial development and hemogenic specification": Table S5

| Table S5 Regulon module activities in VEC clusters, related with Figure 5 | | | | | | | | |
| --- | --- | --- | --- | --- | --- | --- | --- | --- |
|  | EP0 | EP1 | EP2 | EP3 | EP4 | EP5 | EP6 | EP7 |
| Module 1 | -0.895878781 | -0.919503695 | -0.900436536 | -0.848711848 | -1.040417286 | -0.788197847 | -0.800981681 | -0.451215581 |
| Module 10 | -1.654947662 | -1.530108637 | -1.717729701 | -1.583181405 | -1.004391334 | -1.302651194 | -1.88777986 | -2.065970506 |
| Module 2 | 0.066427986 | -0.214817722 | 0.282369731 | 1.354997144 | 0.596891716 | 2.010516529 | -0.295216281 | 0.097834737 |
| Module 3 | 0.108182244 | 0.295482997 | -0.101621919 | -0.396147869 | -0.173579014 | -0.509155912 | 0.867102344 | 0.275637423 |
| Module 4 | 0.943042964 | 1.105849635 | 1.0586645 | 0.97626478 | 2.282082662 | 0.589320562 | 0.958242298 | 0.65108963 |
| Module 5 | -0.123639354 | -0.175697939 | -0.230822281 | -0.032922299 | -0.545370949 | -0.04713406 | 0.444060794 | 0.709977221 |
| Module 6 | 1.576041739 | 1.121727713 | 1.696534048 | 0.652091129 | 0.358786531 | -0.023481528 | 0.845497562 | 1.074749741 |
| Module 7 | -0.93530061 | -0.979515645 | -0.875428093 | -0.783730453 | -0.867005645 | -0.501835154 | -1.216779405 | -1.176210002 |
| Module 8 | -0.158214366 | -0.178679452 | 0.296601111 | 1.216095602 | 0.024255605 | 1.18480006 | 0.376613893 | 0.983005384 |
| Module 9 | 1.074285841 | 1.475262746 | 0.491869141 | -0.554754782 | 0.368747714 | -0.612181456 | 0.709240338 | -0.098898047 |
